# Cannabidiol prevents the locomotor sensitization induced by cocaine and caffeine and upregulates genes of extracellular matrix and anti-inflammatory pathways in the nucleus accumbens: a transcriptome-wide analysis

**DOI:** 10.1101/2023.09.28.560030

**Authors:** José Pedro Prieto, Rafael Fort, Guillermo Eastman, Oliver Kaminski, Carlos Ferreiro-Vera, Verónica Sanchez de Medina, Cecilia Scorza, José Roberto Sotelo-Silveira

## Abstract

Cannabidiol (CBD), a non-psychotomimetic phytocannabinoid found in the *Cannabis* plant, has emerged as a potential therapeutic agent for psychostimulant use disorders. In recent work, we demonstrated that CBD is able to attenuate the expression of locomotor sensitization and the enhanced metabolic activity in the nucleus accumbens (NAc) generated by the combination of cocaine and caffeine. CDB interacts directly or indirectly with several molecular targets, so the underlying mechanisms of its beneficial effects are hard to determine. Here we used high-throughput RNA-sequencing in mice’s NAc after a sensitization protocol with combined cocaine plus caffeine and a CBD pre-treatment, to identify the major pathways and genes involved in CBD attenuating behavioral effects. Results indicated that CBD pretreatment consistently reduced both the acquisition and expression of cocaine and caffeine locomotor sensitization. The transcriptome analysis revealed that CBD pre-treatment enriched genes and functional association between proteins mainly related to extracellular matrix (ECM) organization and cell interactions in the NAc. Moreover, the neuroinflammation and BDNF signaling pathways were also influenced by CBD. Some specially enriched genes such as Tnc were identified as interesting specific candidates for follow-up studies. These findings provide valuable and novel insights into molecular mechanisms of CBD putatively associated with a protective effect on psychostimulant actions. A better understanding of the therapeutic targets of CBD can open new avenues for psychostimulant use disorder treatment strategies.

## Introduction

Cocaine is a highly addictive psychostimulant whose dependence remains a substantial public health challenge, with no effective pharmacological therapy approved to date (Ciccarone and Shoptaw, 2022; Jordan et al., 2019). Its repeated consumption is associated with enduring changes in several brain regions, including the nucleus accumbens (NAc), a key integrator of the mesolimbic reward circuit (Volkow et al., 2019). Cocaine has been described to induce widespread transcriptional and biochemical alterations in the NAc, which may ultimately lead to the maladaptive plasticity that characterizes the development of addiction (Nestler and Luscher 2019; Savell et al., 2020). Besides, forensic data demonstrated that seized samples of cocaine are usually sold adulterated, so these effects can be influenced by the presence of active adulterants, like caffeine (Abin-Carriquiry et al., 2018; Broséus et al., 2016, Cole et al., 2011).

In previous studies, we have demonstrated that caffeine can potentiate several behavioral and molecular actions of cocaine. We have shown that the co-administration of caffeine and cocaine induces a higher acute motor stimulation (López-Hill et al., 2011; Prieto et al., 2012), drug-seeking behavior (Prieto et al., 2016), and wake-promoting effect (Schwarzkopf et al. 2018) than cocaine alone. Caffeine also enhanced the cocaine reward–associated learning, and this was accompanied with an increased expression of immediate-early genes (Muñiz et al., 2017). Additionally, caffeine was able to accelerate and potentiate the expression of cocaine locomotor sensitization, and this response was accompanied by an increased expression of genes related to long-term drug-induced plasticity in the NAc, usually observed after longer, chronic cocaine treatments (Prieto et al., 2015; Prieto et al., 2020a). The importance of the sensitization phenomenon lies in the fact that it reflects persistent neuroadaptations similar to those characteristics of substance use disorders, which underlies the drug-seeking behavior and reinstatement (Steketee and Kalivas, 2011; Vanderschuren and Kalivas, 2000). Based on these results, we postulated that the combination of cocaine and caffeine can accelerate the neuroplastic events that lead to cocaine dependence (Prieto et al., 2020a).

In the last years cannabidiol (CBD) has generated promising perspectives as a pharmacotherapy for substance use disorders (SUDs) (Chye et al., 2019; Gharbi et al., 2023; Navarrete et al., 2021; Paulus et al., 2022). CBD is the most abundant non-psychotomimetic phytocannabinoid of the plant *Cannabis sativa* L (Echeverry et al., 2021b). It has been extensively studied for its potential therapeutic effects in several psychiatric and neurological disorders (Devinsky et al., 2016; Elsaid et al.; 2019; Karimi-Haghighi et al., 2020; Pisanti et al., 2017), and neurodegenerative diseases (Campos et al., 2016; Peres et al. 2018), without t exhibiting large adverse effects (Taylor et al., 2018). Preclinical research on CBD and psychostimulants shows that CBD can exert some protective effects, as it reduces cocaine-induced locomotion (Chesworth and Karl, 2020), drug-seeking behavior (Galaj et al., 2020; Luján et al., 2020), and facilitates the extinction of cocaine conditioned place preference (Calpe-López et al., 2021) in both mice and rats (for a recent review of CBD effects on psychostimulant-induced behaviors see Karimi-Haghighi et al., 2022). However, other studies have revealed contradictory results (Gerdeman et al., 2008; Mahmud et al., 2017; Luján et al., 2018), so the therapeutic potential of CBD on SUDs still needs to be addressed.

Recently, we have shown that CBD was able to attenuate the expression of the locomotor sensitization induced by the combination of cocaine and caffeine, together with the associated metabolic changes found in the rat NAc (Prieto et al., 2020b). However, the underlying mechanisms were not identified since CBD has a multi-target pharmacological profile. CBD interacts directly and indirectly with different neurotransmission systems (i.e., dopaminergic, serotoninergic, endocannabinoid and, opioid systems), signaling pathways (such as BDNF, mTOR and Wnt pathways), and inflammatory response regulators (PPARɤ, IL-1β, IL-6, TNF-α), among others targets (Calpe-López et al., 2019; Karimi-Haghighi et al., 2020; Karimi-Haghighi et al., 2022; Echeverry et al., 2023). Therefore, the present study was designed to identify and investigate the main mechanisms underlying the CBD’s ability to prevent the locomotor sensitization induced by the combined administration of cocaine and caffeine using a genome-wide approximation. To do so, we performed high-throughput RNA-sequencing (RNA-seq) to recognize the major **pathways** and genes in mice’s NAc that were differentially involved in the CBD pre-treatment effects on the cocaine and caffeine sensitization at the transcriptional level. We demonstrated the participation of genes involved in the extracellular matrix (ECM) organization and anti-inflammatory pathways in CBD effects, and revealed novel targets implicated in CBD’s beneficial effects on psychostimulants abuse.

## Materials and methods

### Animals

Male C57BL/6 mice (10–12 weeks old) from the Institut Pasteur of Montevideo animal facilities were employed. All animals were housed in a light- and temperature-controlled room with a 12-h light/dark cycle (lights on at 7:00 am) and had free access to food and water. All experimental procedures were conducted in agreement with the National Animal Care Law (No. 18611) and with the “Guide to the care and use of laboratory animals” (8th edition, National Academy Press, Washington DC, 2010). Furthermore, the Institutional Animal Care Committee approved the experimental procedures. Adequate measures were taken to minimize pain, discomfort or stress of the animals, and all efforts were made to use the minimal number of animals necessary to obtain reliable scientific data.

### Drugs and doses

Cocaine hydrochloride was generously donated by Verardo & Cía Laboratory (Argentina), and caffeine was purchased from Sigma-Aldrich (Germany). Both cocaine and caffeine were dissolved in sterile saline. CBD was kindly donated by Phytoplant Research S.L.U (Spain) and diluted in 3% Tween 80 in sterile saline (vehicle). Cocaine and caffeine were intraperitoneally (i.p.) co-administered at 5 mg/kg and 2.5 mg/kg respectively. CBD was i.p. injected at 20 mg/kg. These doses were selected based on previously published data regarding the facilitating effects of caffeine on cocaine locomotor sensitization (López Hill et al., 2011; Prieto et al., 2015; Prieto et al., 2020a), and the attenuating effect by CBD (Prieto et al., 2020b).

### Behavioral assay: locomotor sensitization

Mice were moved to an experimental room with controlled conditions 72 h previous to the beginning of the experiments. Measurement of locomotor sensitization was carried out using the open field (OF) paradigm, consisting of 30 × 40 × 35 cm acrylic boxes. The animal behavior was recorded and analyzed with the Ethovision XT12 video-tracking software (Noldus, the Netherlands). We measured the horizontal locomotor activity, defined as the total distance moved in meters (m).

The experimental schedule was based on previous studies (Prieto et al., 2020a; Prieto et al., 2020b), and it is shown in Figure 1. One day preceding drug treatment, animals were habituated to the OF for 60 min (day 0), in which basal locomotor activity was recorded. Then, animals were pretreated for 3 days, once a day. Animals received an injection of CBD or its respective vehicle (Veh), and 30 min later were injected with the combination of cocaine and caffeine (CC) or its respective control (Saline). Distance moved in the OF was assessed each day for 60 min after the second injection. After that, animals were kept in their home cages for a 5-day withdrawal period. On day 9, all mice received a challenge dose of CC, and locomotor activity was recorded for 60 min. On days 1 and 9, before the second injection, animals were placed into the OF for a 20-min habituation period (**Figure 1**).

**Figure 1.**
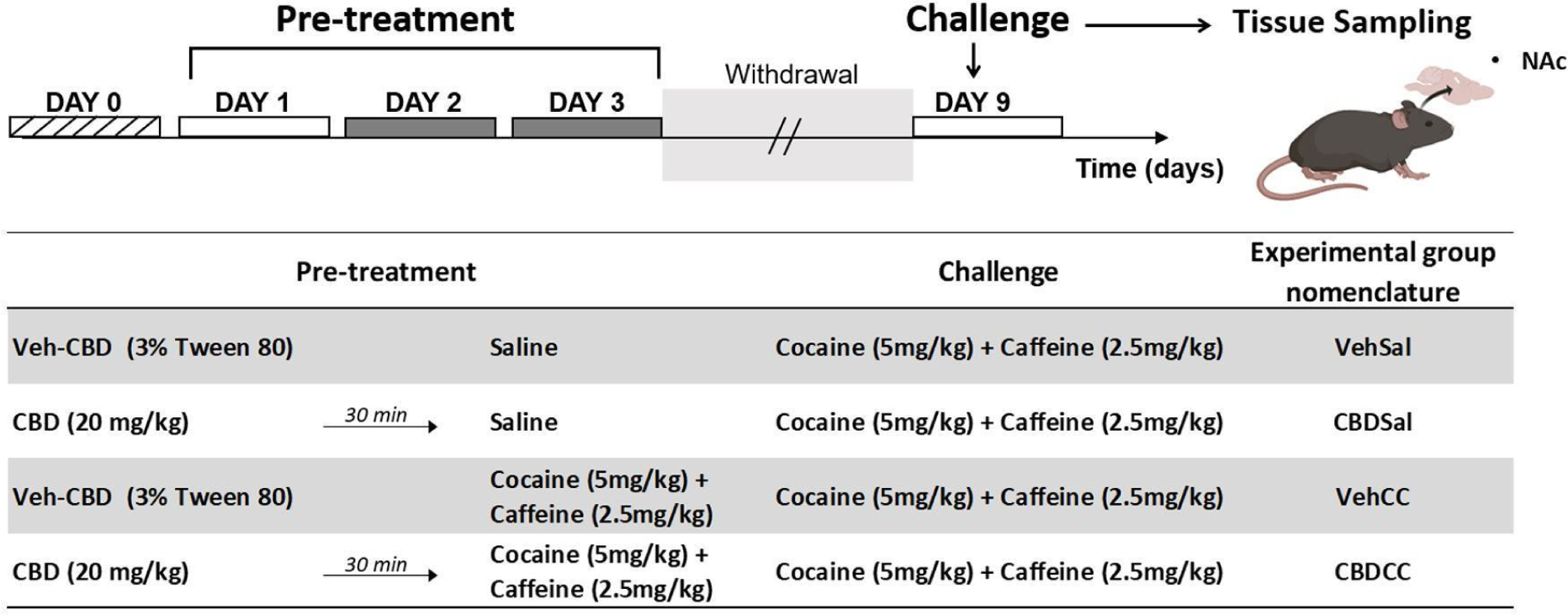
Schematic representation of the experimental protocol used, including the detailed experimental groups, treatment, and nomenclature. Mice were pretreated with CBD or its vehicle (Veh-CBD, 3% Tween 80), and 30 min later received an injection of cocaine + caffeine or saline for 3 days. Five days later (withdrawal period), animals were challenged (day 9) with a dose of cocaine + caffeine. Tissue samples were taken immediately after the end of the behavioral testing on the challenge day (day 9). Modified from Prieto et al. 2020b.

During the experiments, the OF was cleaned with alcohol 30% after each session, before placing the following animal.

### RNA isolation

Brains of male mice were collected immediately after the end of the behavioral session on day 9. NAc punches were taken from individual mice (at least three animals/treatment group) and were rapidly isolated and frozen in RNAlater. The tissue was fully homogenized in 2 ml of QIAzol buffer reagent (Qiagen) on ice and Dounce homogenization. The RNA was extracted using the RNAeasy micro Kit (Qiagen) according to the manufacturer’s instructions. All RNA samples were determined to have 260/280 nm and 260/230 nm absorbance ratio values ≥ 1.8.

### RNA-seq library and sequencing

RNA-seq was performed on samples from individual animals (n ≥ 3) of each experimental group (Figure 1 and Supplementary Figure 1). RNA purity and integrity were assessed using an Agilent 2100 Bioanalyzer System using the RNA 6000 Nano assay (Agilent Technologies). All samples had a RIN value above 7 and libraries were prepared and sequenced by Genewiz (https://www.genewiz.com/) using the TruSeq mRNA Sample Prep Kit v2 protocol (poly(A)+ selection) and Illumina HiSeq 2×150 bp sequencing. The raw FASTQ data sets supporting the results of this article are available at the Sequence Read Archive repository (https://www.ncbi.nlm.nih.gov/sra) (BioProject ID: XXXXXXXXXX).

### RNA-seq analysis

Raw reads (Fastq files) obtained from Genewiz (https://www.genewiz.com/) were used as the input for miARma-Seq pipeline, a comprehensive tool for miRNA and mRNA analysis (Andrés-Leon et al., 2016), following the user manual (v 1.7.2). Low quality reads and adapter sequences were removed with Cutadapt software (Martin, 2011) allowing a minimum read Phred quality of 20. Filtered high quality reads (Ave. 20,059,262 ± SD 2,223,762 reads, **Supplementary Table 1**) were aligned to Mus musculus reference genome (GRCm38/mm10 indexed from http://bowtie-bio.sourceforge.net/bowtie2/index.shtml) using Hisat2 (Kim et al., 2019) with default parameters (Overall genome mapping: Ave. 90.1 ± SD 0.6, **Supplementary Table 1**). Gene counts were assessed with featureCounts (Liao et al., 2014) using default parameters (**Supplementary Table 2**). For quantification we use the annotation coordinates of ensembl Mus_musculus.GRCm38.96 GTF file. Normalization and Differential Expression Genes (DEGs) analysis were conducted using SARTools pipeline (Varet et al., 2016) in RStudio, selecting edgeR algorithm (Robinson et al., 2010), Trimmed Mean of the M-values (TMM) normalization and CPM (Counts Per Million) ≥ 1 as cut-off (**Supplementary Figure 2, Supplementary Table 3 and 4**). A significance threshold of fold change > |1.25| and False discovery rate (FDR) <0.05 or p-value <0.05 was applied for these exploratory studies to retain a sufficient number of genes for conducting over-representation analyses for biological and canonical molecular pathways.

### Gene Ontology, biological process and pathway enrichment analysis

Selected genes were run for over-representation analyses of biological terms and pathways utilizing Enrichr a database that integrates biological data and functional annotation tools (https://maayanlab.cloud/Enrichr/ and database Bioplanet 2019) (Chen et al., 2013). Protein-Protein Interaction analysis (PPIs) was performed using STRING (https://string-db.org/) (Szklarczyk et al., 2019) which integrates functional associations between proteins in a list of genes. Gene Set Enrichment Analysis (GSEA) was performed using gseGO (R package clusterProfiler v4.8.2; Wu et al., 2021) to explore the Gene Ontology (GO) terms (Biological Process, Molecular Function and Cellular Component) using the complete list of genes (ranked based on its fold change of expression), extending beyond the analysis of Differentially Expressed Genes (DEGs). The terms, pathways or networks with significance threshold of p-value <0.05 were selected.

### Hierarchical clustering

Hierarchical clustering was performed with Morpheus software using euclidean algorithm. For the study of cluster of genes modulated by different treatments five range categories (Classes: 2, 1, 0, -1 and -2) were generated and all DEGs were classified in the classes depending of the FC between each comparisons: 2 for DEGs with FC (> 1.5), 1 for DEGs with FC (1.5 > FC > 1.25), 0 for DEGs with FC (1.25 > FC > 0.75), -1 for DEGs with FC (0.75 > FC > 0.5) and -2 for DEGs with FC (<0.5) and hierarchical clustering using euclidean algorithm was performed.

### Statistical analysis

Figures were designed using R and all the analyses were performed in R software (version 3.6) using libraries ggcorrplot, enrichplot, corrplot, xlsx, ggplot2, heatmap.2 (clustering distance measured by Euclidean and Ward clustering algorithms) and Statistica 10 software.

## Data availability

The datasets supporting the conclusions of this article are available at SRA repository (BIOPROJECT) and within the article’s supplementary information.

## Results

### CBD-induced effect on locomotor sensitization elicited by the combination of cocaine and caffeine

First, we analyzed the locomotor activity during the 3 days pretreatment with (1) CBD’s vehicle + saline (VehSal), (2) CBD + saline (CBDSal), (3) CBD’s vehicle + cocaine and caffeine (VehCC), and (4) CBD + cocaine and caffeine (CBDCC). As shown in (**Figure 2A**), 2-way repeated measures ANOVA revealed a significant effect of the treatment [*F*_(3,22)_ = 71.91, p < 0.0001], time [*F*_(2,44)_ = 58.31, p-value < 0.0001] and the treatment × time interaction [*F*_(6,44)_ = 12.23, p-value < 0.0001]. Post hoc analysis showed a gradual and sustained significant increase in the distance moved of the VehCC-treated animals (p-value < 0.001) compared to control group (VehSal) across the 3 days pretreatment. CBDCC group also exhibited a significant increment in its locomotor activity (p-value < 0.001), but limited to days 2 and 3. CBDCC response, however, was lower than VehCC in each day (p-value < 0.05), indicating that CBD administration attenuated the locomotor effects induced by the combination of psychostimulants during the pretreatment phase (**Figure 2A**). There were no significant changes in total locomotor activity between day 1 and day 3 in VehSal or CBDSal-treated animals. On the challenge day, illustrated in (**Figure 2B**), all animals received a dose of cocaine + caffeine. One-way ANOVA revealed a significant effect of treatment [*F*_(3,22)_ = 8.45, p-value < 0.001], and Tukey post hoc test demonstrated a higher locomotor activity of VehCC group compared to VehSal (p-value < 0.05), indicating the expression on locomotor sensitization. CBDCC locomotion, on the other hand, did not differentiate from either the VehSal (p-value = 0.96) nor CBDSal (p-value = 0.17) groups. Furthermore, its activity was significantly lower than the locomotion observed in VehCC group (p-value < 0.05), suggesting that the pretreatment with CBD prevented the expression of locomotor sensitization in the CBDCC group (**Figure 2B**).

**Figure 2.**
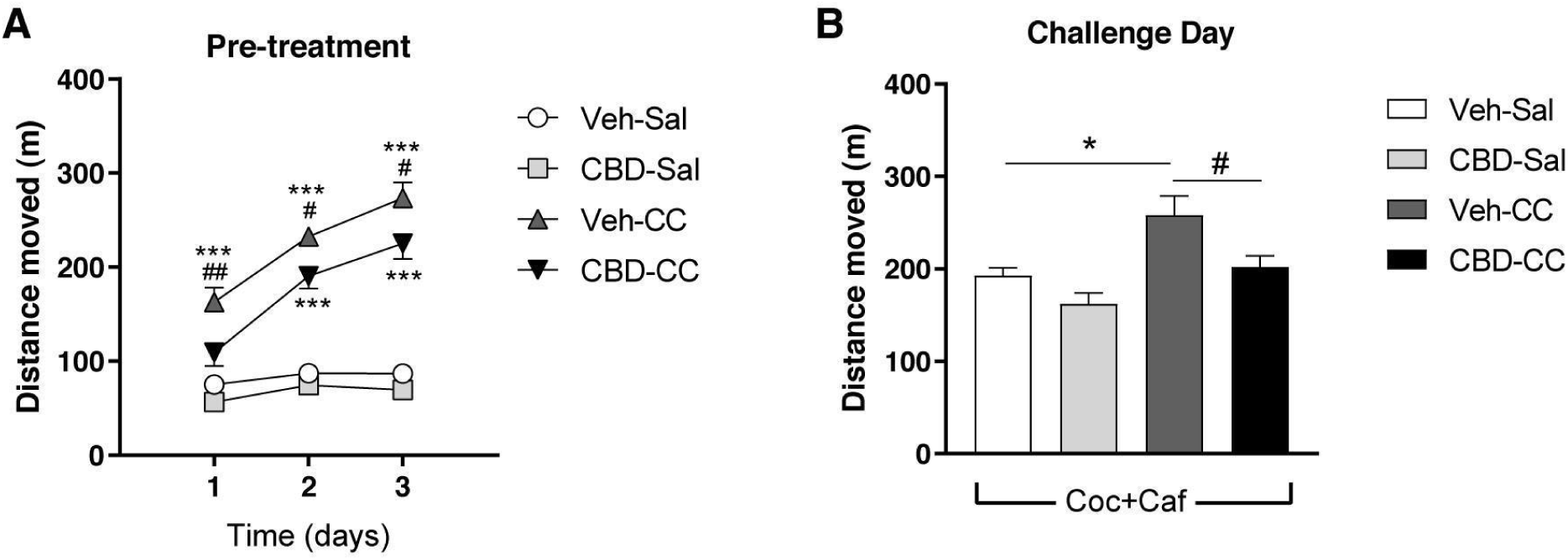
Effect of CBD pretreatment on the locomotor sensitization induced by cocaine + caffeine (CC) during (**A**) pretreatment period, and (**B**) challenge day. Data are expressed as mean ± SEM. Two-way repeated measures ANOVA and one-way ANOVA followed by Tukey’s post hoc test. *p < 0.05 and ***p < 0.001 different from Veh-Sal; #p < 0.05 and ##p < 0.01 different from CBD-CC. N = 6–7.

### Differentially expressed genes (DEGs) and pathways affected by CBD pretreatment

We performed a RNA-Seq of extracted RNA from NAc of mice from each experimental group, immediately after the end of the behavioral session on day 9 (challenge day). A total of approximately 20 million paired-end reads were obtained per sample with an average mapping statistics of 90% aligning to genome and an average of 15 million reads uniquely assigned to genes (see in detail in the method section, **Supplementary Table 1** and **Supplementary Figure 1**).

We first applied the FDR < 0.05 cut-off and fold change > |1.25| to select DEGs and further the analysis. As shown in **Figure 3A**, we identified 17 DEGs, including: *Tnc* (tenascin C)*, Acta2* (actin alpha 2)*, Thbs1* (thrombospondin 1) and *Col5a2* (collagen, type V, alpha 2), that were exclusively upregulated in the groups that received CBD. As for the rest of the genes, *Tnnt1* (troponin T1) and genes from the *Neurod* (neurogenic differentiation 1) and *Slc* (Solute carrier) families were downregulated only in the CBDCC group, while *Entpd4b* (ectonucleoside triphosphate diphosphohydrolase 4B) showed high levels of expression solely in the VehCC group. When we performed the pathway analysis revealed a statistically significant enrichment for terms related to the ECM organization (**Figure 3B**).

**Figure 3.**
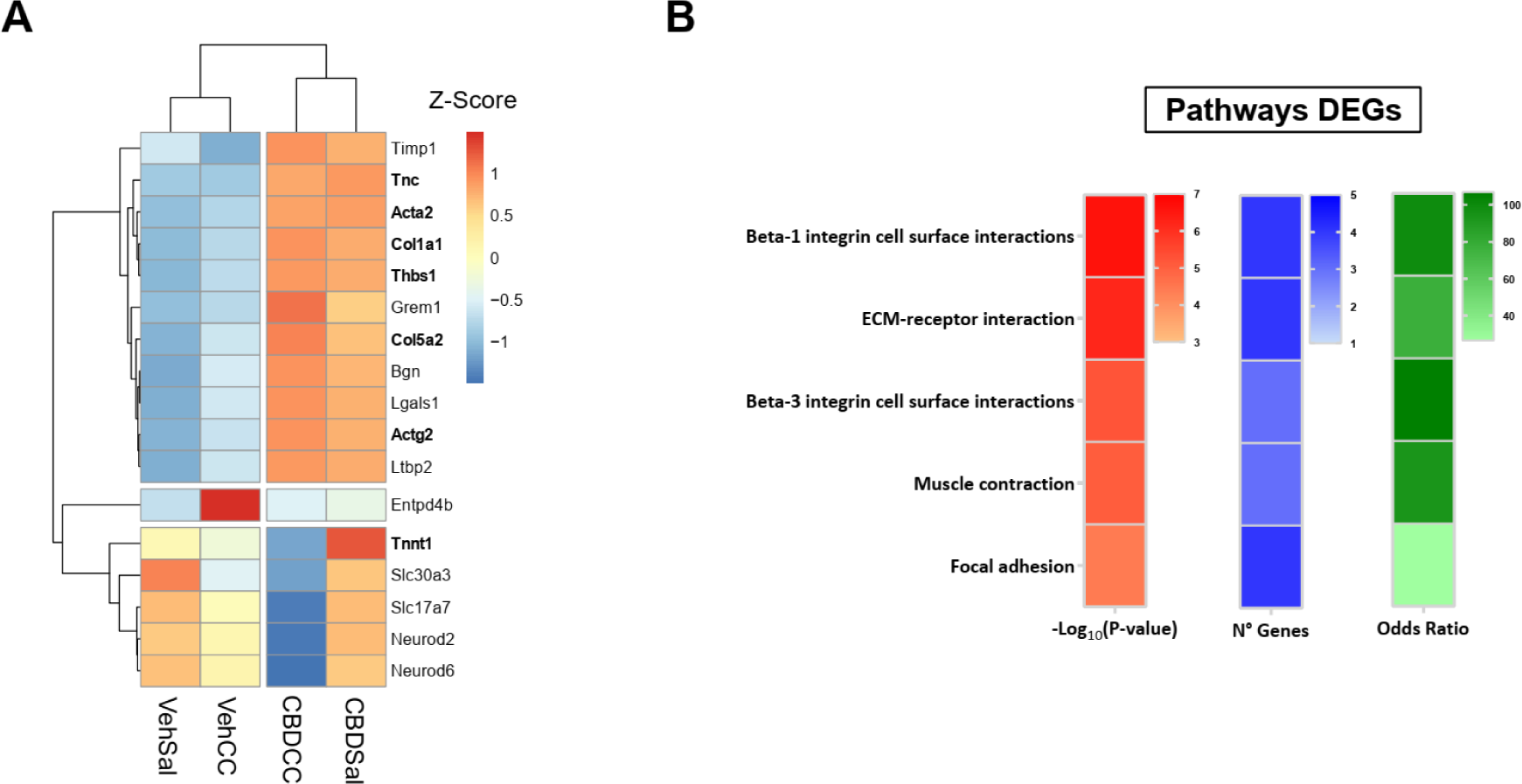
Differentially expressed genes clustering and the enriched pathways and terms associated. A. Hierarchical clustering using euclidean algorithm was performed for DEGs (FDR < 0.05) with Morpheus software. The heatmap of the Z-score for average values groups is shown. In bold are highlighted genes that are enriched in terms and categories shown in B. The top 5 significant terms of the enrichment analysis of pathways and processes among the DEGs are shown, heatmaps in red, blue and green expresses -log10(p-value), number of genes found associated in the term/category and odd ratio, respectively.

Then we used a broader DEG selection to retain a sufficient number of genes for conducting over-representation analyses for biological and canonical molecular pathways (as done previously in related works; Farris et al., 2015; Lo Iacono et al., 2016; Powell et al., 2020; Walker et al., 2018). We identified a broader list of DEGs (p-value<0.05 and fold change>|1.25|) of each treatment versus the control group (VehSal) (**Figures 4A-C** and **Supplementary Table 4**) and the DEGs of CBDCC versus VehCC (**Figure 4D**). Interestingly, the only comparison without CBD (see **Figure 4A**), involving the sensitized group versus VehSal, showed the weaker transcriptional response, with 120 DEGs (see **Supplementary Table 4**). It is noteworthy, however, the strong upregulation of the *Entpd4b* gene in VehCC (**Figure 4A**). All the groups that received CBD shared the significant upregulated genes, such as *Acta2*, *Thbs1*, *Lgals1* (lectin, galactose binding, soluble 1), *Col5a2* and *Tnc* (**Figure 4B-D**), and both CBDSal versus VehSal, and CBDCC vs VehCC showed a moderately robust transcriptional response (320 and 284 DEGs, respectively). When compared to the sensitized group, CBDCC also observed an important downregulation of *Entpd4b* (**Figure 4D**).

**Figure 4.**
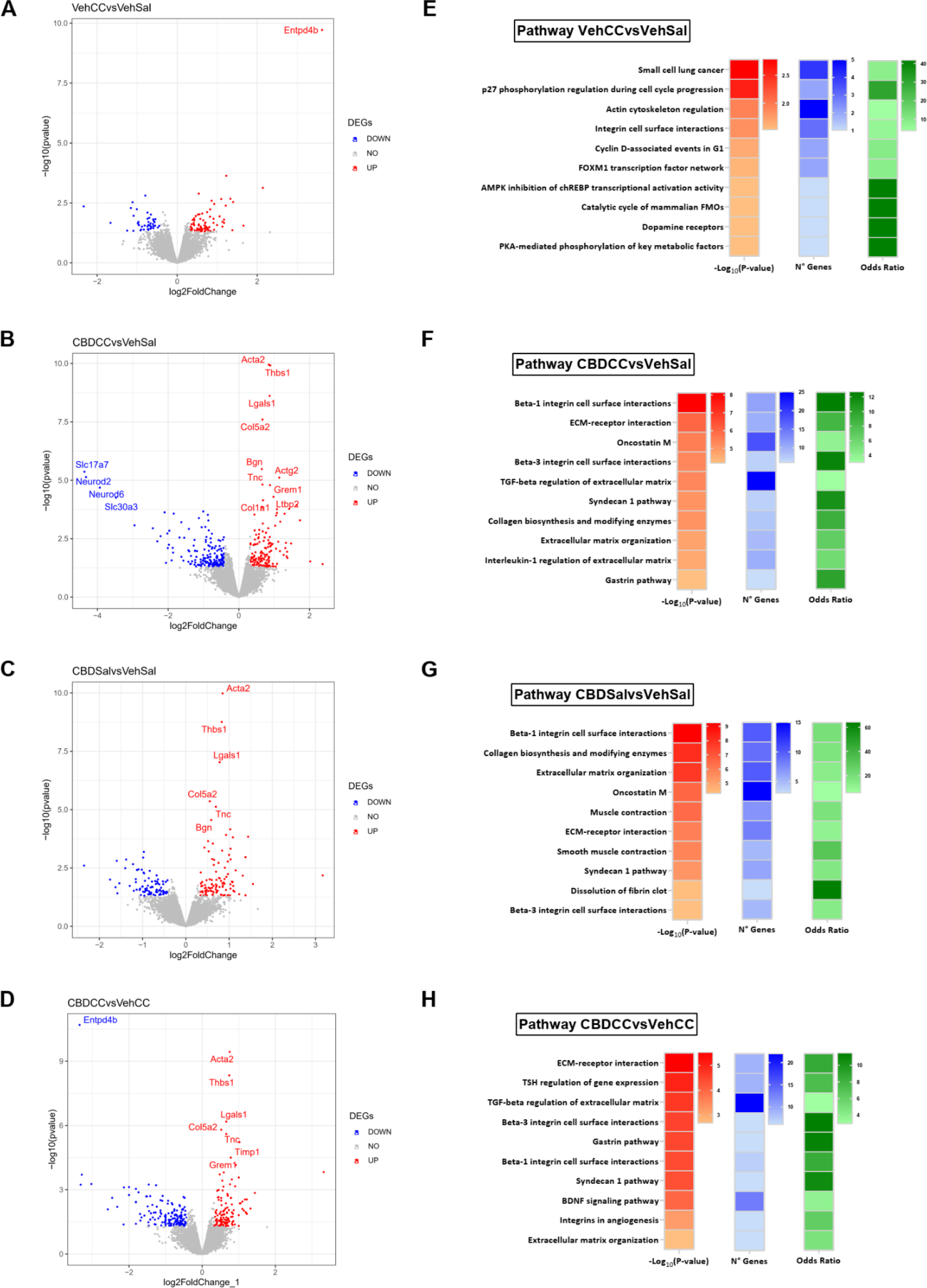
Genes differentially expressed, pathways and terms associated with the different groups comparison. Volcano plots are shown, the abscissa shows the relative change values expressed as log2(FC) and the ordinate shows the p-values associated expressed as -log10(p-value). In red and blue are highlighted genes upregulated or downregulated, respectively with a p-value < 0.05 and fold change > |1.25|; and the label name is shown for those genes with FDR < 0.05. The top 10 significant terms of the enrichment analysis of pathways and processes among the DEGs are shown, heatmaps in red, blue and green expresses -log10(p-value), number of genes found associated in the term/category and odd ratio, respectively. **A** and **E**. The CBD’s vehicle + cocaine and caffeine (VehCC) vs CBD’s vehicle + saline (VehSal) comparison is shown. **B** and **F**. The CBD + cocaine and caffeine (CBDCC) vs CBD’s vehicle + saline (VehSal) comparison is shown. **C** and **G** The CBD + saline (CBDSal) vs CBD’s vehicle + saline (VehSal) comparison is shown. **D** and **H**. The CBD + cocaine and caffeine (CBDCC) vs CBD’s vehicle + cocaine and caffeine (VehCC) comparison is shown.

To explore the functions of the identified DEGs list, we analyzed which pathways and cellular processes were enriched for each comparison using Enrichr (https://maayanlab.cloud/Enrichr/ and the database Bioplanet 2019) (Chen et al., 2013) (see **Supplementary Table 5**). Among the top ten significant pathways enriched in the VehCC group, we found the dopamine receptors pathway; pathways involved in energy metabolism such as AMPK inhibition of chREBP transcriptional activation activity and PKA-mediated phosphorylation of key metabolic factors; and actin cytoskeleton regulation pathway (**Figure 4E**). The groups that were pretreated with CBD exhibited a significant enrichment of several pathways involved in ECM organization, including ECM-receptor interaction, interleukin-1 regulation of ECM, collagen biosynthesis and modifying enzymes, TGF-beta regulation of ECM, ECM organization, and integrin cell surface interactions (**Figure 4F-H**), Of interest regarding the psychostimulant actions, in the CBDCC versus VehCC comparison, the BDNF signaling pathway was also enriched.

To surpass the constraints of the over-representation analysis based on a limited number of DEGs, we used GSEA analysis to confirm the pathways seen above using an alternative type of calculation involving the whole set of genes in the transcriptome. This well-known strategy explores the complete list of genes ranked based on the fold change of expression and conducts an enrichment analysis of Gene Ontology terms using the ClusterProfiler algorithm (Wu et al., 2021). This approach showed similar enrichment terms (**Figure 5, Supplementary Table 6 and Supplementary Figure 3**) when compared to the terms defined by DEGs approach in **Figure 4**. Particularly noticeable are the positive enriched terms related to ECM organization after CBD pretreatment, in contrast to the observed suppression of microtubule and dynein motility-related terms (**Figure 5 and Supplementary Figure 3**).

**Figure 5.**
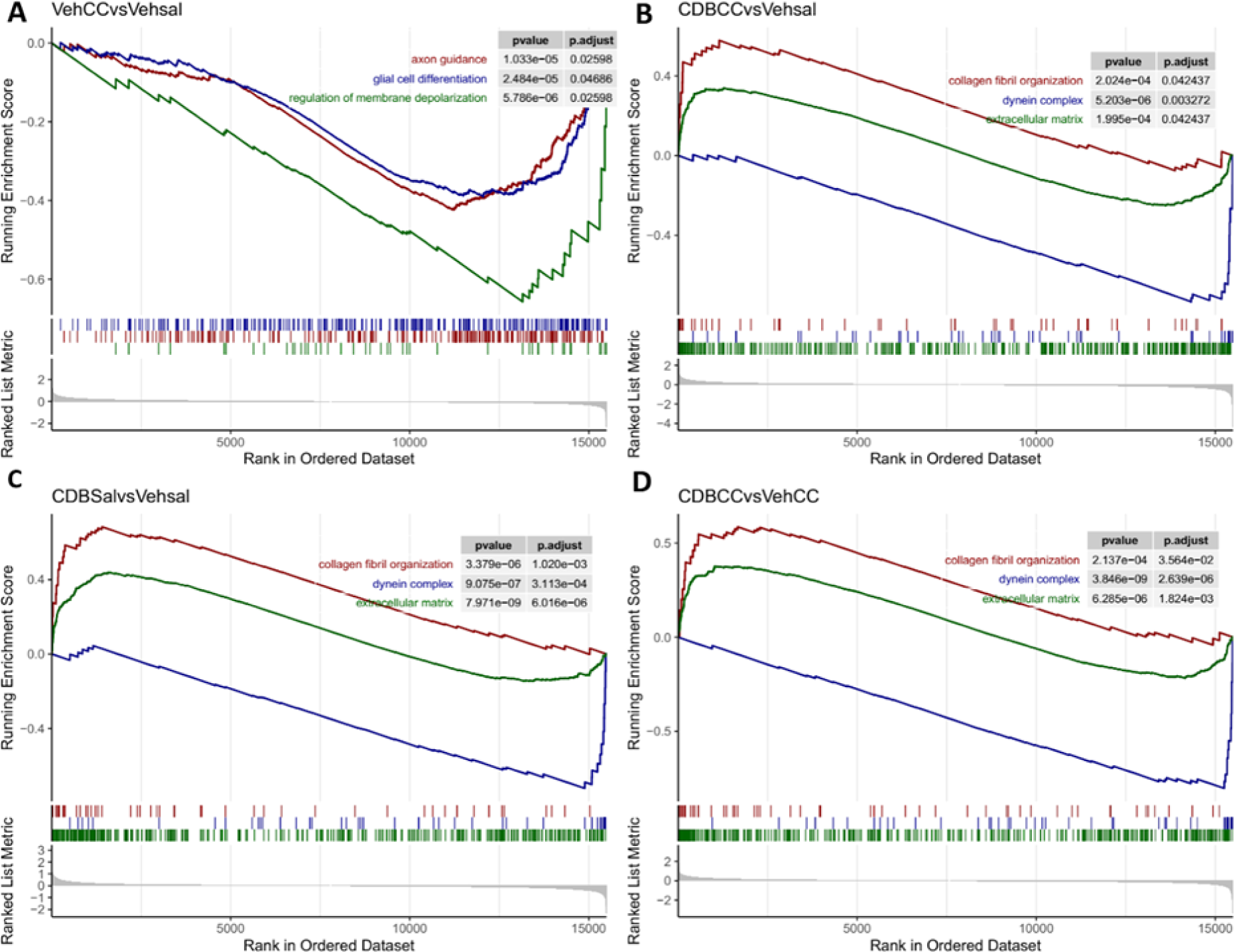
Gene Set Enrichment Analysis (GSEA) of Gene Ontology terms associated with the different groups comparison. GSEA plot of three selected statistically significant enriched GO terms for each of the comparisons. The x-axis represents the rank for all genes; the y-axis (upper panel) shows the Enrichment Score value, the y-axis (bottom panel) represents the value of the ranking metric. **A**. The CBD’s vehicle + cocaine and caffeine (VehCC) vs CBD’s vehicle + saline (VehSal) comparison is shown. **B**. The CBD + cocaine and caffeine (CBDCC) vs CBD’s vehicle + saline (VehSal) comparison is shown. **C**. The CBD + saline (CBDSal) vs CBD’s vehicle + saline (VehSal) comparison is shown. **D**. The CBD + cocaine and caffeine (CBDCC) vs CBD’s vehicle + cocaine and caffeine (VehCC) comparison is shown.

In order to have a comprehensive view of the different responses induced by the CBD pretreatment, we also compared the enrichment and odds ratio of selected pathways and biological processes between the previous comparisons, based either on their consistency or the direct relation with cocaine and caffeine mechanism of action. The result of this analysis is shown in **Figure 6**. The pathways related with the ECM organization showed the highest levels of enrichment, present almost entirely in the comparisons that included the CBD pretreatment. Globally, these results evidence an important action of CBD over the organization of the ECM. A similar pattern was observed for the BDNF signaling pathway. On the other hand, only the comparisons with VehCC showed an enriched dopamine receptors pathway, with higher representation in the sensitized group. Moreover, only the sensitized group was associated with the enrichment of both dopamine receptors and caffeine metabolism pathways, in accordance with the molecular actions of cocaine and caffeine.

**Figure 6.**
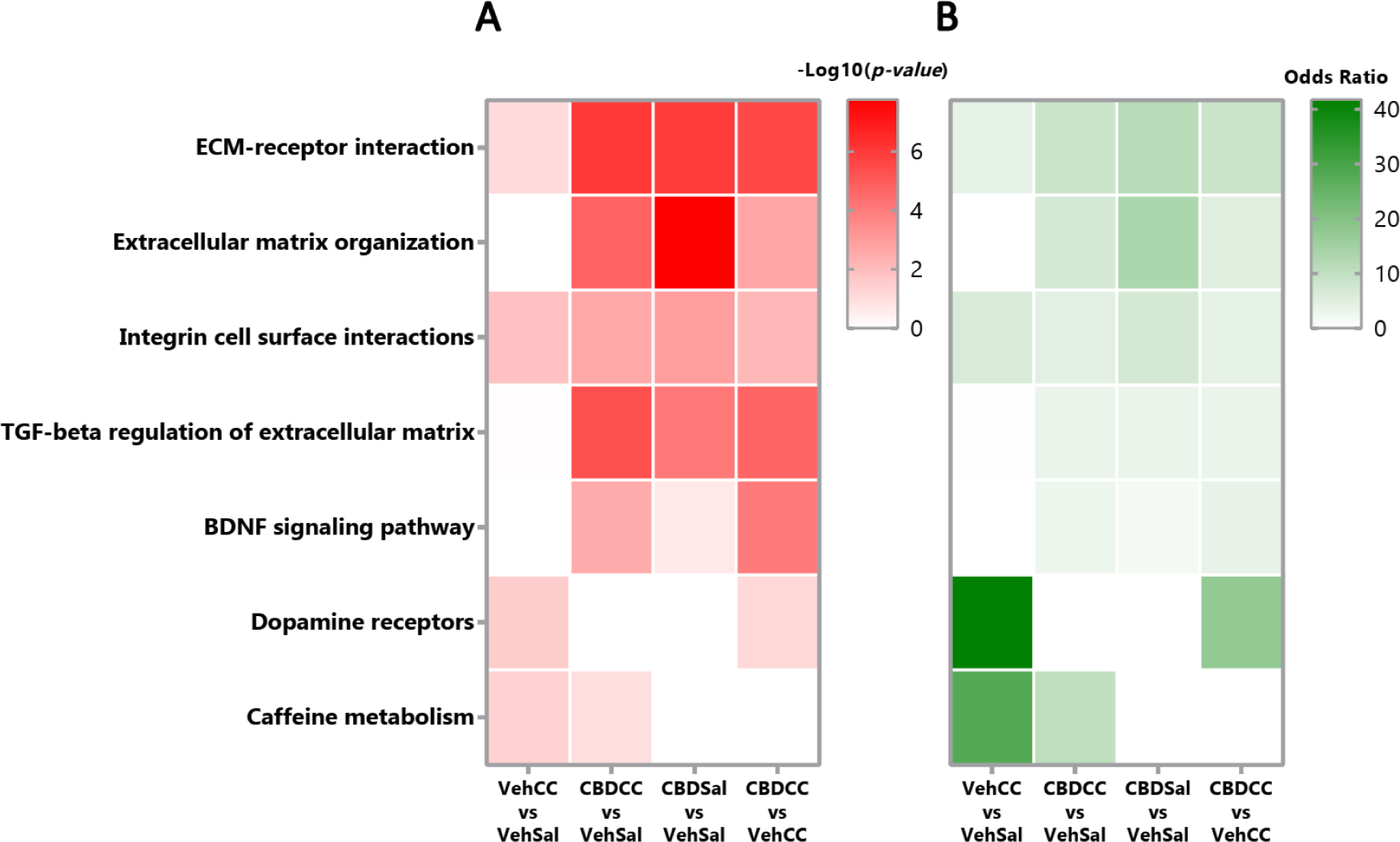
Selected pathways and terms for the different group comparisons are shown. A and B. Heatmaps of the -log10(p-value) and Odds ratio respectively, for the different comparisons evaluated.

### Candidate genes involved in the effects of CBD

We were then interested in identifying a comprehensive list of genes whose expression could be modulated in clusters among the treatment conditions. For that purpose, we classified genes among conditions in different classes and clusterized (**Figure 7**). This approach led us to identify 3 clusters based on one of the following conditions: (1) mRNAs had a peak of expression only in the sensitized group; (2) or a peak of their expression with CBD pretreatment; and (3) bottomed their expression with CBD pretreatment. The gene names included in these clusters and their relative change in expression induced by the treatments compared to the control group (VehSal) are shown (**Figure 7B-D**). We next used the list of genes obtained to perform a protein-protein interaction analysis of each cluster. Functional associations between proteins were present solely in cluster 3, containing those genes that increased their expression in the groups that received CBD but not in the sensitized one (**Figure 7D**). The protein interaction network included several collagen alpha chains and Thbs1, among others. Most of these genes were enriched in the ECM pathway, and some of them were also enriched in integrin cell surface interaction and ECM-receptor interaction pathways; again, indicating a prominent role of the ECM and its cell interaction in CBD effects. In the case of Tnc, although it did not show protein associations, it was enriched in all three named pathways, suggesting a relevant involvement in the observed response.

**Figure 7.**
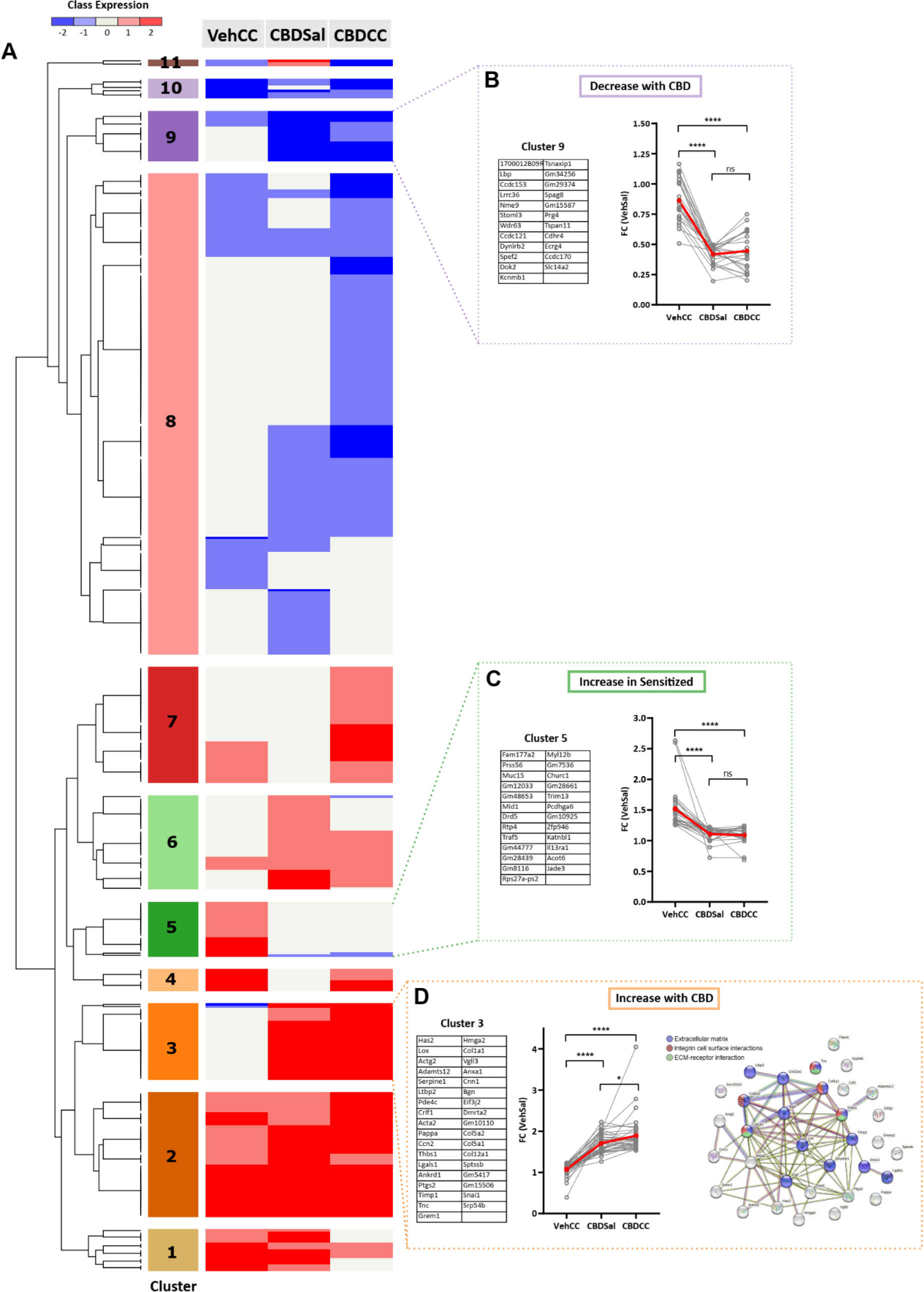
Cluster of genes modulated by different treatments. A. Hierarchical clustering using euclidean algorithm was performed for categorized DEGs with Morpheus software. Five range categories (Classes) were generated and all DEGs were classified depending of the FC between each comparison: 2 for DEGs with FC (> 1.5), 1 for DEGs with FC (1.5 > FC > 1.25), 0 for DEGs with FC (1.25 > FC > 0.75), -1 for DEGs with FC (0.75 > FC > 0.5) and -2 for DEGs with FC (<0.5). Three clusters were selected to show the genes included in them: (B) Cluster 3 (a cluster of genes increased in CBD-treated groups), (C) Cluster 5 (a cluster of genes increased in the Sensitized group) and (D) Cluster 9 (a cluster of genes decreased in CBD treated groups). For the three selected clusters the original FC values are plotted and the gene names are shown. The three selected clusters were analyzed with STRING and only cluster 9 showed an enriched PPIs (p < 0,05) and three significant enriched biological processes (p < 0,05) are highlighted with colors. Two-tailed t-test was applied for comparisons and ns for non-significant; * for p < 0,05 and **** for p < 0,0001 was used.

## Discussion

This study confirms our previous finding that CBD is able to prevent the locomotor sensitization induced by the combination of cocaine and caffeine, and shows that this effect could be associated with a CBD-induced differential increase in the expression of genes related to the interaction and organization of the ECM in the NAc. Our results contribute to the identification of the mechanisms and candidate genes putatively associated with the effects of CBD on psychostimulant sensitization.

Here, we used an established administration protocol that induces a fast acquisition and expression of locomotor sensitization of low doses of combined cocaine and caffeine (Prieto et al., 2015; Prieto et al., 2020a, Prieto et al., 2020b). Previously, we had shown that the sensitized animals with the co-administration of these psychostimulants exhibited increased metabolic activity in the NAc (Prieto et al., 2020b), as well as specific changes in the mRNA expression of receptor subunits of dopaminergic and glutamatergic systems, and some synaptic plasticity associated genes (Prieto et al., 2020a). In the present study, the sensitized group presented an enrichment of pathways and biological terms involved in energy metabolism, dopaminergic receptors and caffeine metabolism, in agreement with our previous observations. As expected, the sensitized animals had a higher representation of genes of the dopaminergic and caffeine metabolism pathways, reflecting a stronger alteration of pathways related to the direct molecular actions of cocaine and caffeine in the group without CBD. Among the DEG’s induced by cocaine and caffeine co-administration, we found that *Entpd4b* was particularly upregulated in the sensitized animals. *Entpd4b* encodes an apyrase involved in nucleotide recycling (Gorelik et al., 2020), which may suggest an increased adenosine production as a compensatory mechanism to the adenosine receptors blockage by caffeine (Fisone et al., 2004). Overall, our results in the NAc of the sensitized animals agree with previous data showing the increased recruitment of the NAc as a relevant component in the expression of psychostimulant-induced behavioral sensitization (Steketee and Kalivas, 2011).

CBD pretreatment was able to both decrease the acquisition and prevent the expression of cocaine and caffeine locomotor sensitization, which reaffirms our previous observations (Prieto et al., 2020b). Given the multiple targets of CBD, a myriad of possible mechanisms has been proposed to underlie its effects on psychostimulant behavioral actions, mostly involving the modulation of neurotransmitter systems, signal transduction pathways and neuroinflammation (Karimi-Haghighi et al., 2022). In this study, all the groups that received CBD showed a consistent enrichment of pathways and biological terms involved with the interaction and organization of the ECM. This included the CBDCC vs VehCC comparison, which indicates a possible involvement of the ECM in the observed effects of CBD on locomotor sensitization. This was an initially surprising effect, as there is not much data regarding the involvement of the ECM in the CBD mechanisms as an addiction-protective substance. The activation of these pathways, however, could reflect the participation of neural plasticity processes. CBD has been shown to modulate both plasticity and cell proliferation in the hippocampus of psychostimulant-treated animals (Luján et al., 2018; Razavi et al., 2021), associated with reduced drug-reinforcement (Luján et al., 2018; Luján et al., 2020). In the NAc, CBD was also able to rescue maladaptive synaptic plasticity induced by a binge of alcohol (Brancato et al., 2022), showing also a modulatory action in this region. Additionally, the ECM could be affected directly by CBD, as it can inhibit the activity of matrix metalloproteinases (MMPs). MMPs catalyze the decomposition of several components of the ECM, such as collagen and elastin, and interact with adhesion molecules such as integrins (Chen et al., 2023; Yue et al., 2012). This action may contribute to the prevention of the neural plasticity mechanisms involved in repeated drug exposure and sensitization. In this sense, our results showed that the genes increased with CBD were enriched in biological pathways related with ECM organization, and integrin and ECM-receptor interaction.

Considering the above-mentioned relation between CBD, ECM and plasticity, it is noteworthy that the groups treated with CBD also showed an increased enrichment of BDNF (Brain Derived Neurotrophic Factor) signaling pathway, which is not present in the sensitized group. BDNF is known to promote neuronal growth, differentiation, synaptogenesis (Lu and Figurov, 1997), and has been extensively studied as an important modulator of the drugs-induced neuroadaptations in the NAc (Bolaños and Nestler, 2004; Ghitza et al., 2010). A BDNF increase in the NAc has been shown to promote drug reinforcement and drug-seeking behavior (Bahi et al., 2008; Graham et al., 2007), but also to inhibit drug-induced plasticity (Anderson et al., 2017). On the other hand, CBD upregulation of BDNF in other areas, such as hippocampus, amygdala and prefrontal cortex, has been proposed as one mechanism by which CBD may reduce psychostimulants reinforcing effects (Calpe-López et al., 2019). Here, we showed that CBD influences the BDNF pathway, mainly in CBDCC compared to the sensitized group, hence suggesting a similar involvement in the NAc. Recently, Shen and collaborators (Shen et al., 2022) showed that CBD modulation of the BDNF-TrkB signaling pathway inhibits the dopamine release induced by methamphetamine. A similar mechanism may be in play here, preventing the consolidation of the neural changes involved in the cocaine and caffeine locomotor sensitization.

The implication of the ECM in the CBD-induced effects is reinforced by the fact that the cluster of genes upregulated by CBD and involved in the ECM and cell receptor interactions, were the only ones that exhibited protein functional associations. Among them, the most significant DEGs in that cluster were *Acta2* (Alpha-Actin-2), *Col5a2* (Collagen Type V Alpha 2 Chain), *Thbs1* (Thrombospondin-1) and *Tnc* (Tenascin-C, TnC). Thrombospondin-1 is an adhesive protein that facilitates cell-ECM interaction, promotes angiogenesis, and can interact with MPPs (Bein and Simons, 2000). TnC, in turn, is an ECM protein that participates in cell adhesion, cell proliferation and migration, and binds to different matrix components, soluble factors and pathogens (Midwood et al., 2016). Interestingly, an increased expression of TnC is observed in inflammation and other pathological conditions (Abedsaeidi et al., 2023). TnC has an active role in promoting both pro- and anti-inflammatory responses (Abedsaeidi et al., 2023; Imanaka-Yoshida et al., 2020; Yalcin et al., 2020), and has been shown to regulate chemotaxis and cytokine production in microglia (Haage et al., 2019). Furthermore, studies using TnC-deficient mice have demonstrated an increased Iba1 immunoreactivity during post-ischemic neuroinflammation (Manrique-Castano et al., 2021). Neuroinflammation is currently considered one of the processes that contribute to the neural adaptations that occur after chronic exposure to drugs of abuse and sensitization (Kohno et al., 2019; Lacagnina et al., 2017). For example, repeated psychostimulant administration has been associated with an increase in microglial Iba1 expression in the striatum (Liao et al., 2016). For its part, the CBD anti-inflammatory property is well established (Atalay et al., 2020), and large piece of evidence indicates that CBD is able to decrease psychostimulant-induced pro-inflammatory responses, such as glial reactivity and cytokine production, among others (Calpe-López et al., 2019; Karimi-Haghighi et al., 2019). Taken together, this data suggests that the anti-inflammatory action of CBD could participate in the attenuating effect of the locomotor sensitization observed in this work. TnC seems to be an interesting candidate to further study the precise mechanisms of CBD to prevent the repeated and endurance action induced by psychostimulant drugs, involving the ECM and inflammatory response.

The present study describes the transcriptional response of the combined cocaine and caffeine locomotor sensitization in the NAc, and identifies mechanisms and pathways —ECM-cell interactions, BDNF and neuroinflammation— involved in CBD’s protective effects on cocaine and caffeine-induced sensitization. The engagement of these pathways by CBD could be explained by the already known neuroprotective effect of CBD described in different contexts (e.g., neurodegeneration) that involve structural and inflammatory changes (di Giacomo et al., 2020; Echeverry et al., 2021a; Martín-Moreno et al., 2011).

Our findings point out specific candidate genes, that given the relevance of the NAc in psychostimulant reinforcement and addiction, may be of interest to investigate further to better dissect CBD therapeutic targets in psychostimulant use disorders.

## Acknowledgments

This study has a license from IRCCA, Uruguay. Cecilia Scorza thanks the Ibero-American Cyted-CannaLatan network.

## Supplementary Figures

**Supplementary Table 1.**
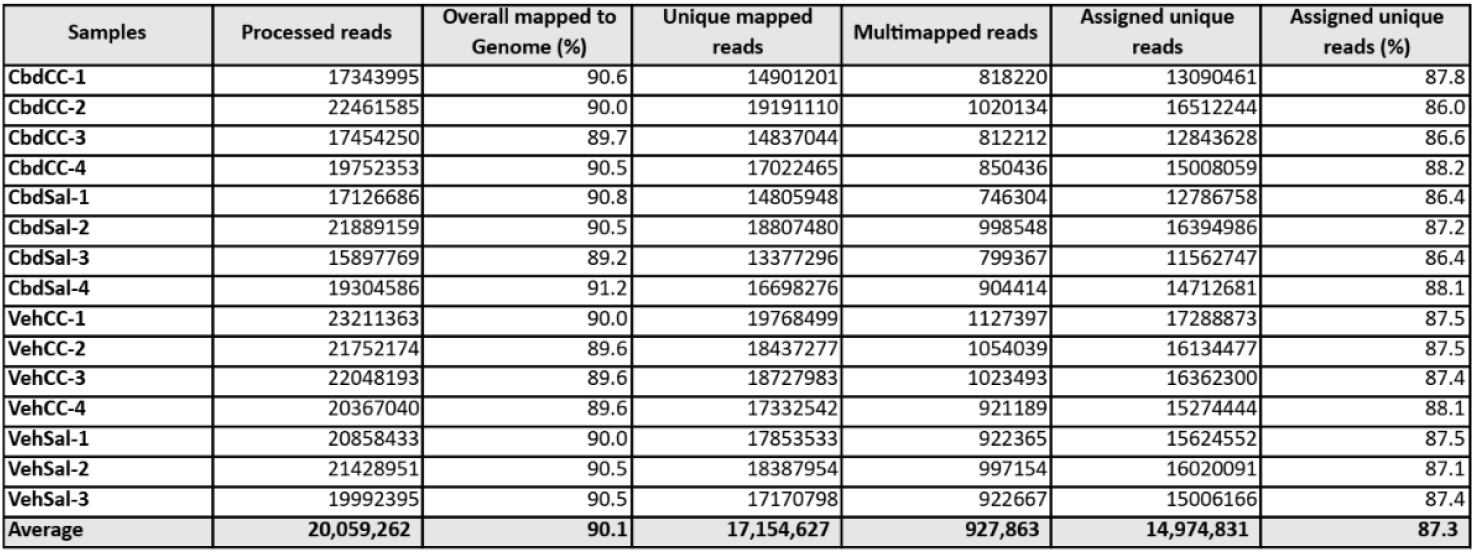

**Supplementary Figure 1.**
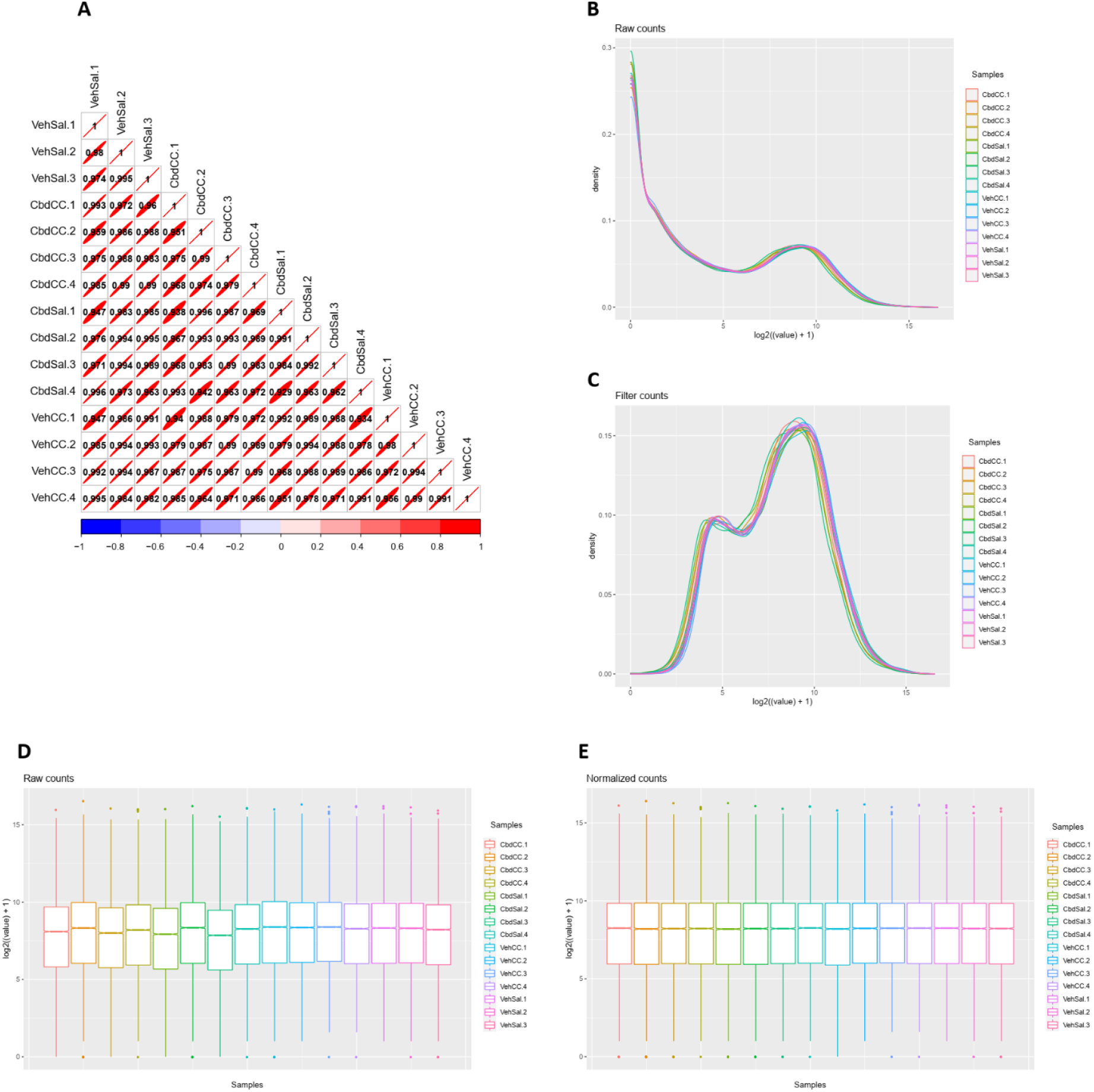
Analysis and processing of the RNA-seq samples. **A**. Pearson correlation matrix of the samples. Density plot of the samples pre (**B**) and post (**C**) filtering low read counts features. Boxplot of the samples pre (**D**) and post (**E**) normalization was applied.

**Supplementary Figure 2.**
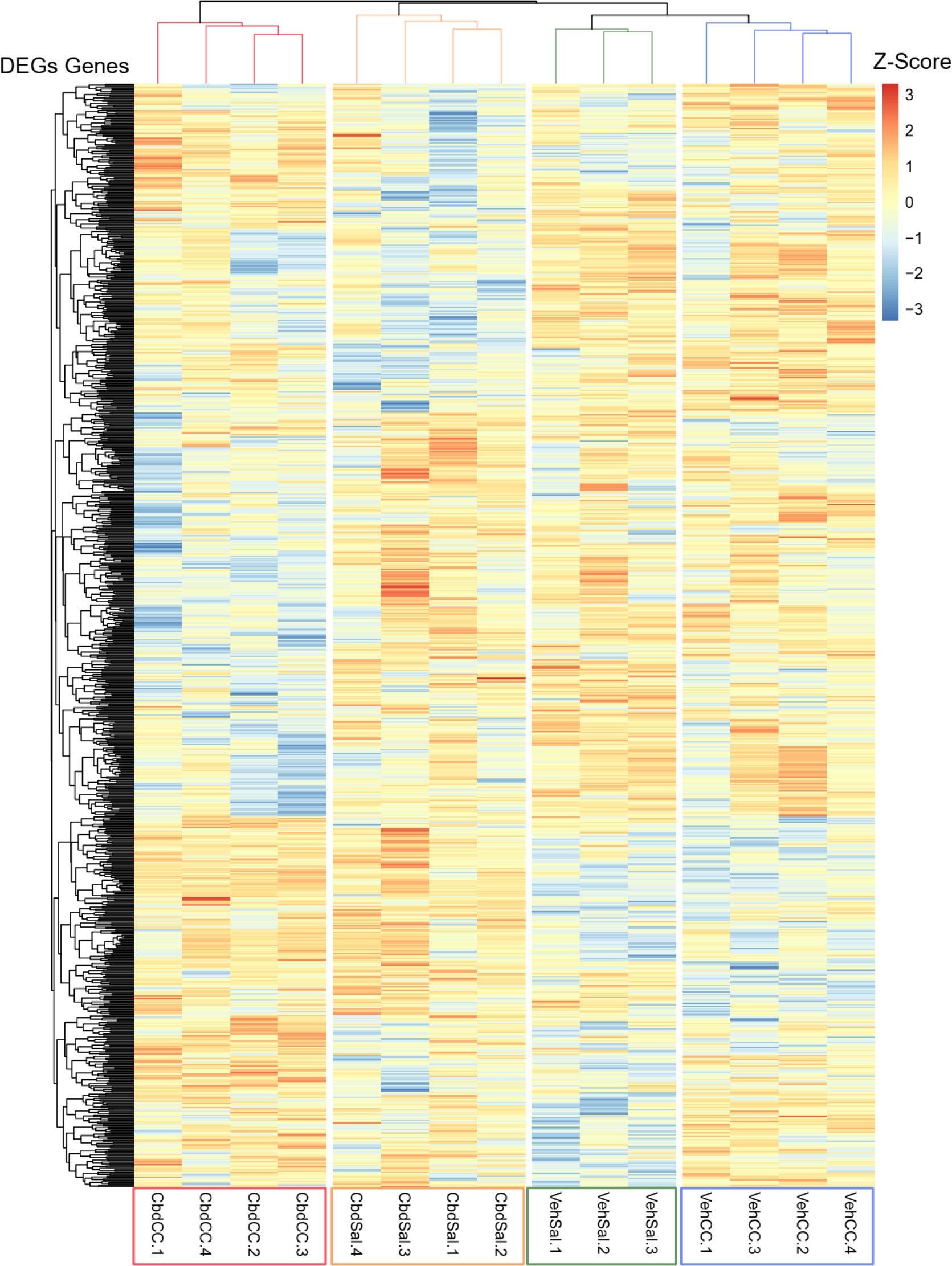
Differentially expressed genes. Heatmap of differentially expressed genes (DEGs, p-value<0.05 and fold change>|1.25|) for the different comparisons, with expression shown as Z-score of log2 normalized counts (two-way hierarchical clustering distance measured by Euclidean and Ward clustering algorithms).

**Supplementary Figure 3.**
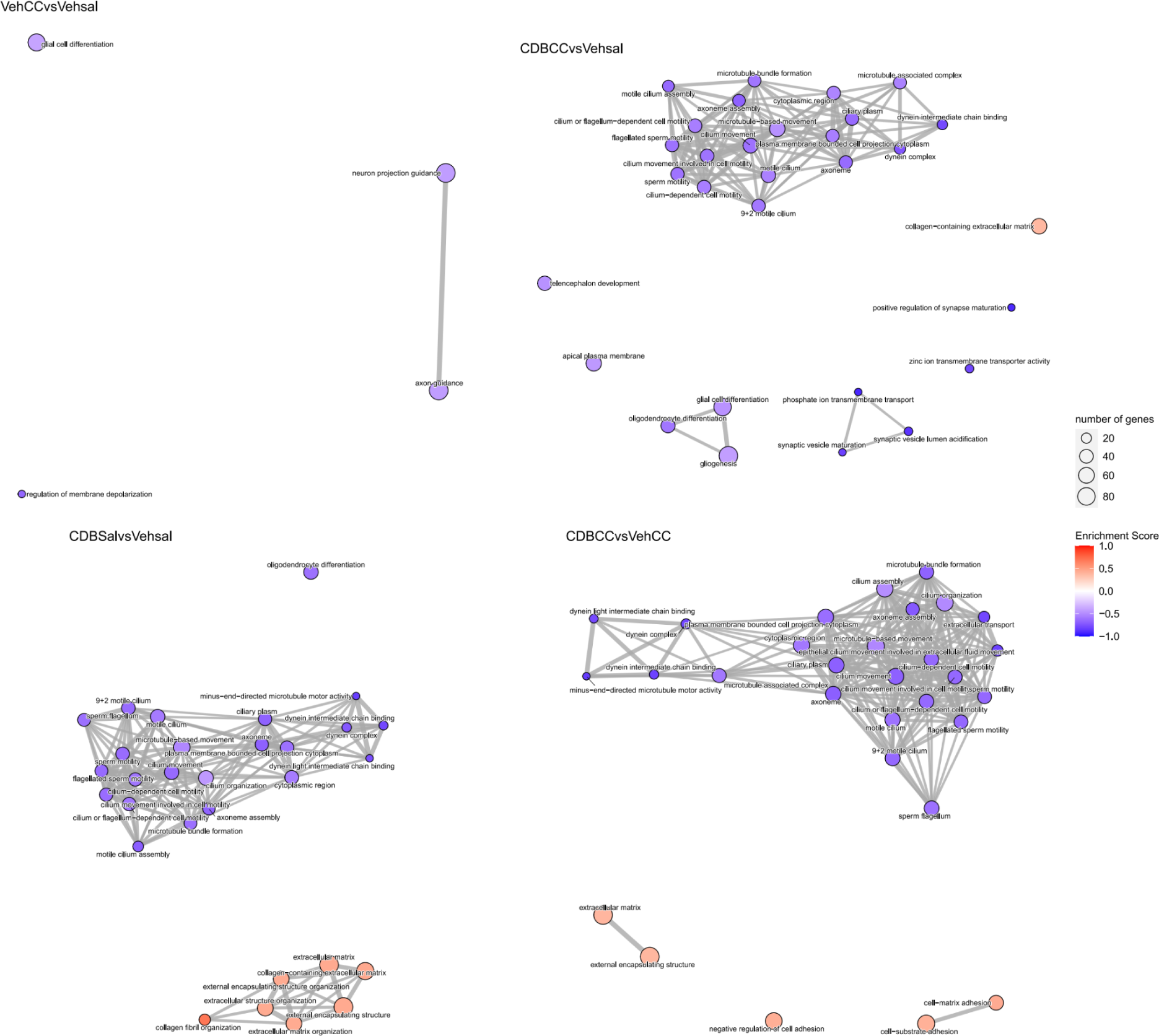
Gene Ontology term functional enrichment with ClusterProfiler for the different comparisons. Emapplot of significantly enrriched GO terms for the different comparisons analyzed. The nodes represent GO terms and the connections represent shared genes, the Enrrichment Score is a statistical measure of the degree of enrichment for the GO term categories within the gene dataset based on the ranked list of genes.

