## Supplemental Figures for "Cannabidiol prevents the locomotor sensitization induced by cocaine and caffeine and upregulates genes of extracellular matrix and anti-inflammatory pathways in the nucleus accumbens: a transcriptome-wide analysis"

### Supplementary Figures:

Supplemetary Table 1

| Samples | Processed reads | Overall mapped to Genome (%) | Unique mapped reads | Multimapped reads | Assigned unique reads | Assigned unique reads (%) |
|---|---|---|---|---|---|---|
| **CbdCC-1** | 17343995 | 90.6 | 14901201 | 818220 | 13090461 | 87.8 |
| **CbdCC-2** | 22461585 | 90.0 | 19191110 | 1020134 | 16512244 | 86.0 |
| **CbdCC-3** | 17454250 | 89.7 | 14837044 | 812212 | 12843628 | 86.6 |
| **CbdCC-4** | 19752353 | 90.5 | 17022465 | 850436 | 15008059 | 88.2 |
| **CbdSal-1** | 17126686 | 90.8 | 14805948 | 746304 | 12786758 | 86.4 |
| **CbdSal-2** | 21889159 | 90.5 | 18807480 | 998548 | 16394986 | 87.2 |
| **CbdSal-3** | 15897769 | 89.2 | 13377296 | 799367 | 11562747 | 86.4 |
| **CbdSal-4** | 19304586 | 91.2 | 16698276 | 904414 | 14712681 | 88.1 |
| **VehCC-1** | 23211363 | 90.0 | 19768499 | 1127397 | 17288873 | 87.5 |
| **VehCC-2** | 21752174 | 89.6 | 18437277 | 1054039 | 16134477 | 87.5 |
| **VehCC-3** | 22048193 | 89.6 | 18727983 | 1023493 | 16362300 | 87.4 |
| **VehCC-4** | 20367040 | 89.6 | 17332542 | 921189 | 15274444 | 88.1 |
| **VehSal-1** | 20858433 | 90.0 | 17853533 | 922365 | 15624552 | 87.5 |
| **VehSal-2** | 21428951 | 90.5 | 18387954 | 997154 | 16020091 | 87.1 |
| **VehSal-3** | 19992395 | 90.5 | 17170798 | 922667 | 15006166 | 87.4 |
| **Average** | **20,059,262** | **90.1** | **17,154,627** | **927,863** | **14,974,831** | **87.3** |

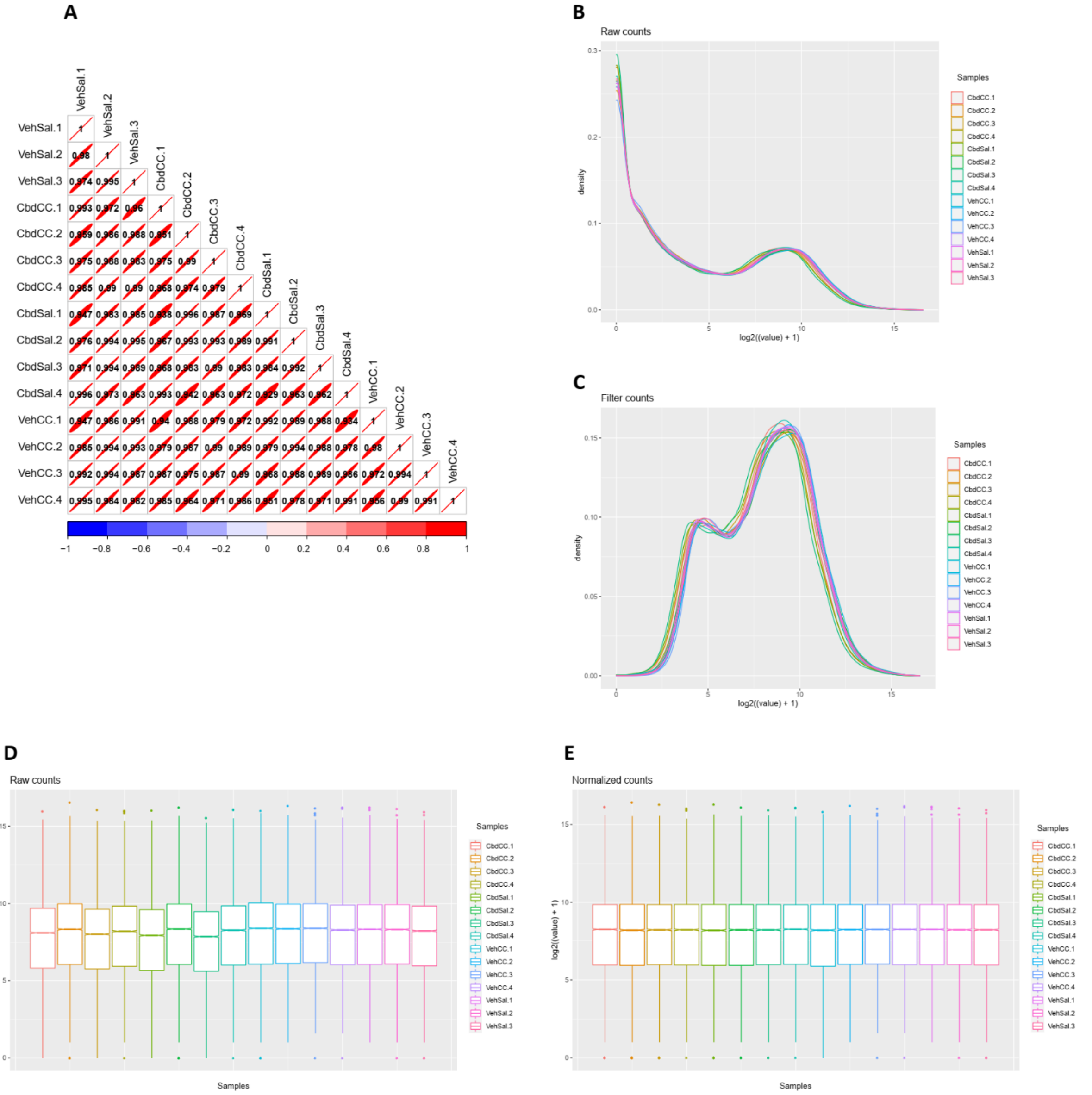


**Supplementary Figure 1.** Analysis and processing of the RNA-seq samples. **A**. Pearson correlation matrix of the samples. Density plot of the samples pre (**B**) and post (**C**) filtering low read counts features. Boxplot of the samples pre (**D**) and post (**E**) normalization was applied.

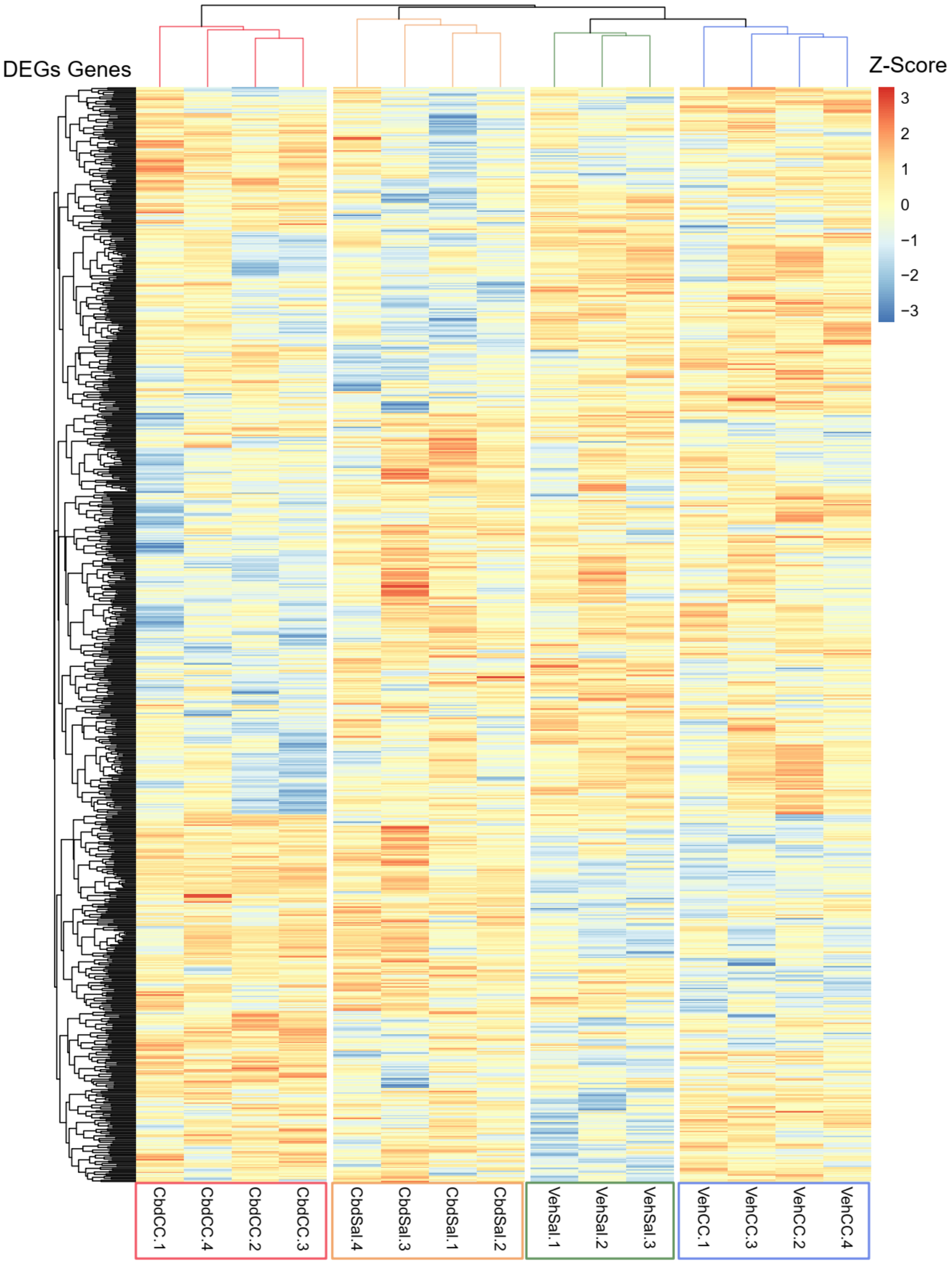


**Supplementary Figure 2.** Differentially expressed genes. Heatmap of differentially expressed genes (DEGs, p-value<0.05 and fold change>|1.25|) for the different comparisons, with expression shown as Z-score of log2 normalized counts (two-way hierarchical clustering distance measured by Euclidean and Ward clustering algorithms).

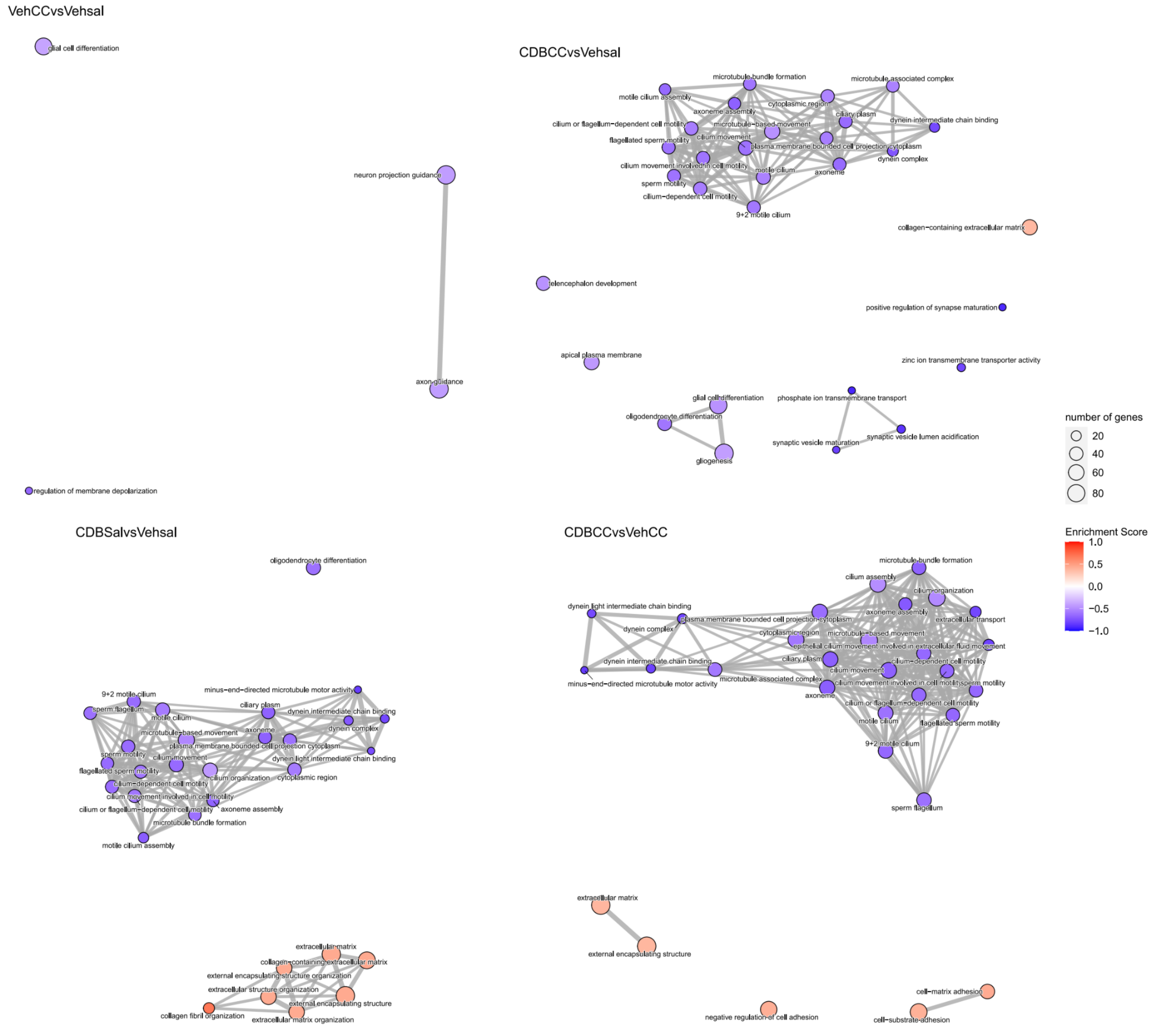


**Supplementary Figure 3.** Gene Ontology term functional enrichment with ClusterProfiler for the different comparisons. Emapplot of significantly enrriched GO terms for the different comparisons analyzed. The nodes represent GO terms and the connections represent shared genes, the Enrrichment Score is a statistical measure of the degree of enrichment for the GO term categories within the gene dataset based on the ranked list of genes.
